# High-throughput profiling of metabolic responses to exogenous nutrients in *Synechocystis* sp. PCC 6803

**DOI:** 10.1101/2023.11.15.567228

**Authors:** Vilhelmiina Haavisto, Zachary Landry, Sammy Pontrelli

## Abstract

Cyanobacteria fix carbon dioxide and release carbon-containing compounds into the wider ecosystem, yet they are sensitive to small metabolites that may impact their growth and physiology. Several cyanobacteria can grow mixotrophically, but we currently lack a molecular understanding of how specific nutrients may alter the compounds they release, limiting our knowledge of how environmental factors might impact primary producers and the ecosystems they support. In this study, we develop a high-throughput phytoplankton culturing platform and identify how the model cyanobacterium *Synechocystis sp.* PCC 6803 responds to nutrient supplementation. We assess growth responses to 32 nutrients at two concentrations, identifying 15 that are utilized mixotrophically. Seven nutrient sources significantly enhance growth, while 19 elicit negative growth responses at one or both concentrations. High-throughput exometabolomics indicates that oxidative stress limits PCC 6803’s growth, but may be alleviated by antioxidant metabolites. Furthermore, glucose and valine induce strong changes in metabolite exudation in a possible effort to correct pathway imbalances or maintain intracellular elemental ratios. This study sheds light on the flexibility and limits of cyanobacterial physiology and metabolism, as well as how primary production and trophic food webs may be modulated by exogenous nutrients.

**Importance:** Cyanobacteria capture and release carbon compounds to fuel microbial food webs, yet we lack a comprehensive understanding of how external nutrients modify their behavior and what they produce. We developed a high throughput culturing platform to evaluate how the model cyanobacterium *Synechocystis sp.* PCC 6803 responds to a broad panel of externally supplied nutrients. We found that growth may be enhanced by metabolites that protect against oxidative stress, and that growth and exudate profiles are altered by metabolites that interfere with central carbon metabolism and elemental ratios. This work contributes a holistic perspective of the versatile response of PCC 6803 to externally supplied nutrients, which may alter carbon flux into the wider ecosystem.

## Introduction

Cyanobacteria are primary producers found in virtually all terrestrial and aquatic ecosystems with access to light, where they carry out essential biogeochemical and ecological functions. Most importantly, they play prominent role in global carbon fixation[1,2], where they form the base of the food web by releasing carbon-containing compounds that support the growth of heterotrophic microbes in the surrounding environment [3,4]. However, cyanobacteria also respond sensitively to exogenous nutrients that may originate from anthropogenic or other biological sources[5] [6,7]. Changes to cyanobacterial physiology may have cascading effects on the composition and behavior of the heterotrophic bacterial community and the wider ecosystem, underscoring the importance of understanding how a broad range of exogenous nutrients affect cyanobacteria and other primary producers on a molecular level.

The effects of certain exogenous nutrients on cyanobacterial growth and population dynamics have been observed previously. For example, in aquatic ecosystems, eutrophication can spur large blooms as cyanobacteria take advantage of excess nitrogen and phosphorus[5], while chemical pollution can restructure entire communities by eliminating sensitive species and selecting for resistant ones[6,7]. Alternatively in the lab, cultures supplemented with carbon sources can either experience growth inhibition or grow mixotrophically, using auto– and heterotrophic modes of metabolism simultaneously[8,9]. Mixotrophy can be an important determinant of microbial fitness in many habitats[10,11]. For example, in aquatic environments, mixotrophic growth may enable the highly abundant marine cyanobacterium *Prochlorococcus* to survive at depths where light penetration is insufficient for them to survive on photoautotrophic growth alone[12].

Though growth and population-level responses have been characterised, we still lack a comprehensive understanding of how exogenous nutrients influence cyanobacterial exudation, which may have cascading effects as exuded compounds fuel heterotrophic microbes. These microbes not only carry out functions that are important for the broader ecosystem [13–16] [17] [18], but also engage in a bidirectional relationships with phototrophs, as they produce metabolic byproducts and micronutrients [19] or remineralized high molecular weight photosynthate[13,20] that can, in turn, be used mixotrophically by cyanobacteria.

Cyanobacterial mixotrophic capabilities have thus far been demonstrated across phylogenetically disparate species, using carbon sources including acetate[21,22], glycerol[8], and some amino acids[9] and monosaccharides[8]. Yet, cyanobacteria encode a wide range of transporters for compounds including amino acids and oligopeptides[23] and urea and sugars[24], suggesting that the range of mixotrophically assimilable compounds may be wider than what has been shown for any individual species so far.

Here, we explore the mixotrophic capabilities of the model cyanobacterium *Synechocystis sp.* PCC 6803. We determine how supplementation of a broad range of metabolites alters its growth and exudate to understand the physiological processes that may be perturbed by exogenous nutrients. To overcome the inherent throughput limitations of typical, flask-based cyanobacterial cultivation, we developed a high throughput cyanobacterial culturing platform to systematically screen the growth and exudation behaviour of PCC 6803 supplemented with 32 metabolites at two different concentrations. We measured nutrient uptake and exudation using high-throughput exometabolomics, allowing us to identify cellular processes that define how PCC 6803 responds to exogenous nutrients.

## Results

### Development of a high-throughput cultivation system

Evaluating cyanobacterial responses to a wide range of growth conditions has historically been limited by cultivation throughput. Cyanobacteria, as well as other photoautotrophic microorganisms, are commonly cultured in flask-based setups[25–27] that are inherently low-throughput. However, microtiter plates that are routinely used in high-throughput bacterial and fungal cultivation present key challenges for the successful cultivation of photoautotrophic organisms including cyanobacteria. In particular, sufficient gas exchange must be possible to avoid carbon dioxide (CO_2_) limitation, which reduces photoautotrophic growth[28]. CO_2_ limitation is easily overcome in flasks by sparging in ambient air or CO_2_[26,29], but this solution is not possible to implement with microtiter plates.

There are several other options to mitigate CO_2_ limitation in microtiter plates, but each has critical disadvantages. For example, ‘gas-permeable’ microplate seals often prevent both gas exchange and evaporation[30] and can also prevent light from reaching the cells. Besides inadequate gas exchange and potential shading, microplate seals are also impractical for work that requires regular access to the cultures for sampling. CO_2_ can also be supplemented by adding sodium bicarbonate directly to the culture medium[31]; however, this can lead to alkalinisation as the bicarbonate buffer equilibrates with atmospheric CO_2_, which can in turn inhibit cyanobacterial growth[32].

One possibility to overcome pH changes caused by bicarbonate supplementation is to buffer the medium. To test the feasibility of this approach, we performed an abiotic incubation of BG-11 medium supplemented with various concentrations of sodium bicarbonate and HEPES buffer. We observed that even low concentrations of sodium bicarbonate (10 mM) could not be buffered by the highest tested concentration of HEPES (50 mM) to a pH below 8 **(Fig. 1A).** Sodium bicarbonate is usually supplied at concentrations between 10-50 mM [33–35], and by decreasing the concentration below 10 mM to reduce alkalinization, we would likely lose its positive effect on growth. Meanwhile, further increasing HEPES concentrations could induce osmotic changes that would alter cyanobacterial physiology, which is also undesirable. Considering these limitations, we concluded that bicarbonate supplementation is not appropriate for our high-throughput culturing platform.

**Figure 1:**
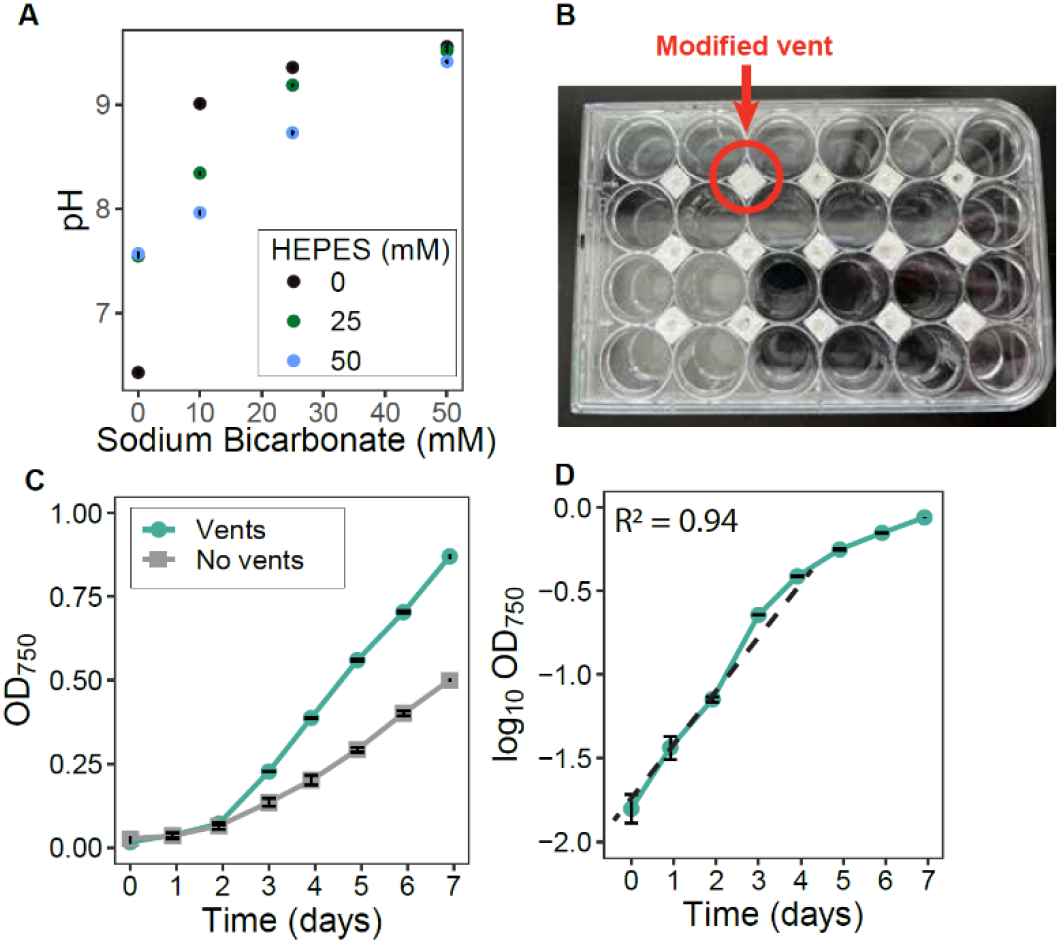
Growth of PCC 6803 in an optimized high-throughput cultivation setup. All error bars represent standard deviation around the mean. A) Abiotic incubation of different concentrations of HEPES buffer and sodium bicarbonate in BG-11 medium. B) Photograph of the modified 24-well plate lid on top of a commercially available 24-well plate. C) OD_750_ growth curves of PCC 6803 in 24-well plates with and without vented lids. D) The vented lids allow PCC 6803 to grow exponentially in the first 4 days. The dashed line indicates a linear regression between day 0 and 4, with the R squared indicated on the plot. All error bars represent standard deviation of the mean of three independent cultures.

To overcome CO_2_ limitation, we modified a microtiter plate lid, placing vents between the wells for aeration, which were covered with a PVDF membrane (pore size 0.45 µm) to prevent airborne contamination **(Fig. 1B)**. These lids allowed us to achieve robust cyanobacterial growth in 24-well plates while relying on atmospheric CO_2_ concentrations **(Fig. 1C, D)**, with exponential growth occurring in the first four days. We chose to use 24-well plates to maximize the volume available for sampling over longer experiments and to improve aeration in the wells. The plates also allowed us to measure cyanobacterial growth *in situ* using a plate reader. Altogether, we developed and implemented a culturing platform that allowed us to expand the number of different growth conditions we can test at once.

### Growth response to metabolite supplementation

Using our high-throughput culturing platform, we tested how nutrient supplementation influences the growth of *Synechocystis* sp. PCC 6803. We grew PCC 6803 under a 12:12 light:dark cycle and in the presence of one of 32 metabolites that represent a broad range of amino acids, organic acids, monosaccharides and C1 compounds. We included compounds that have not previously been tested for consumption by PCC 6803 during growth, and glucose and acetate which have widely been recognized as mixotrophic substrates for PCC 6803[36,37]. Of the chosen compounds, several have previously been shown to be incorporated into PCC 6803 biomass using ^14^C labelled substrates (Table 1). However, these experiments typically measure uptake within seconds to minutes, which do not reflect whether the metabolite inhibits growth, can be used as a source of carbon or nitrogen to substantially improve biomass yield, or alters PCC 6803’s exudation behaviour. We supplemented metabolites at a high concentration of 5 mM and a lower concentration of 100 μM, yielding a total of 65 growth conditions including the non-supplemented control. These concentrations were chosen to broadly cover the large range of metabolite concentrations that have been supplemented to PCC 6803 or other cyanobacteria in previous works[8,9,22,38]. We monitored growth using OD_750_ **(Fig. 2A)** as a proxy for cyanobacterial abundance, and measured metabolite consumption after ten days of growth using liquid chromatography quadrupole time of flight mass spectrometry (LC-QTOF-MS).

**Figure 2:**
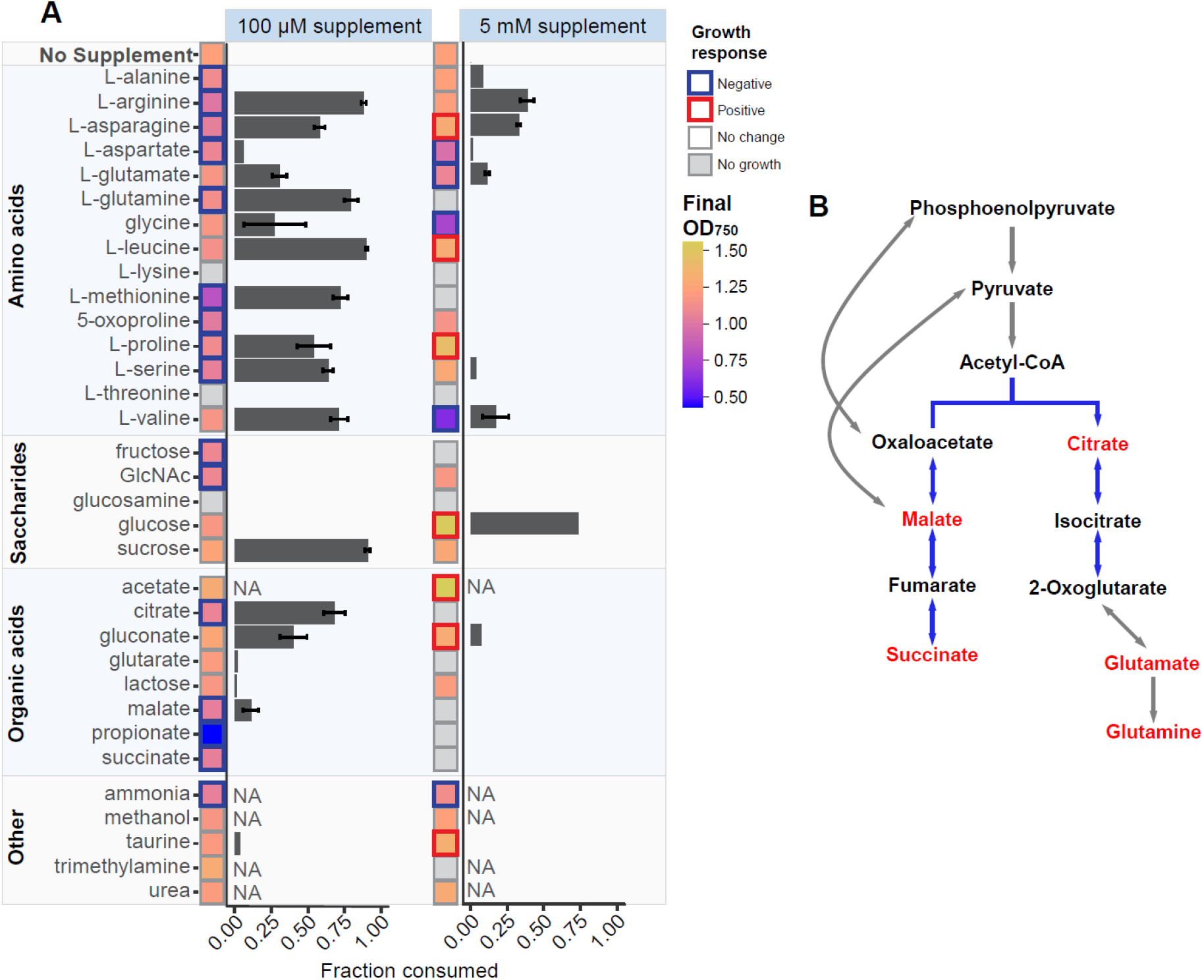
Growth and metabolite consumption. A) Growth of PCC 6803 after 10 days with supplementation of metabolites at 100 μM or 5 mM concentrations. Grey squares signify conditions where no growth is observed, and colours represent final OD_750_ after 10 days. The border represents conditions where there is a significant (p < 0.05, Welch’s t test) increase (red) or decrease (blue) in OD_750_ compared to the no supplement control. Horizontal bars represent the fraction of the supplemented metabolite that is consumed after 10 days of growth. Compounds whose fraction consumed is marked “NA” are unable to be measured using the applied metabolomics method. All experiments were performed in triplicate, and error bars represent standard deviation around the mean. B) Diagram of the branched TCA cycle in PCC 6803 (blue lines) and related reactions. Metabolites in red inhibit growth when supplemented.

**Table 1:** Metabolites that PCC 6803 is known to consume, observed either in this or previous works. ^14^ C incorporation refers to uptake of isotopically labelled metabolites by PCC 6803, measured in seconds to minutes. Medium drawdown refers to measurements of metabolite depletion in culture medium of PCC 6803 after growth.

| Compound | Method | Reference |
| --- | --- | --- |
| *Sugars* |  |  |
| Glucose | Medium drawdown | [36] This study |
| Sucrose | Medium drawdown | This study |
| *Organic acids* |  |  |
| Gluconate | Medium drawdown | This study |
| Malate | Medium drawdown | This study |
| Citrate | Medium drawdown | This study |
| Acetate | Medium drawdown | [37] |
| *C1 compounds* |  |  |
| Urea | C<sup>14</sup> incorporation | [73] |
| *Amino acids* |  |  |
| L-Alanine | C<sup>14</sup> incorporation | [38] |
| L-Arginine | C<sup>14</sup> incorporation | [38,74] |
|  | Medium drawdown | This study |
| L-Asparagine | C<sup>14</sup> incorporation | [38] |
|  | Medium drawdown | This study |
| L-Glutamate | C<sup>14</sup> incorporation | [38] |
|  | Medium drawdown | This study |
| L-Glutamine | C<sup>14</sup> incorporation | [38,74] |
|  | Medium drawdown | This study |
| Glycine | C<sup>14</sup> incorporation | [38] |
|  | Medium drawdown | This study |
| L-Histidine | C<sup>14</sup> incorporation | [38,74] |
| L-Leucine | C<sup>14</sup> incorporation | [38] |
|  | Medium drawdown | This study |
| L-Lysine | C<sup>14</sup> incorporation | [38,74] |
| L-Methionine | Medium drawdown | This study |
| L-Phenylalanine | C<sup>14</sup> incorporation | [38,74] |
| L-Proline | C<sup>14</sup> incorporation | [38] |
|  | Medium drawdown | This study |
| L-Serine | C<sup>14</sup> incorporation | [38] |
|  | Medium drawdown | This study |
| L-Valine | Medium drawdown | This study |

Of the 64 conditions with metabolite supplementation, seven significantly improved the growth of PCC 6803 (p < 0.05, Welch’s t-test), while 36 partially or completely inhibited growth. These growth effects were highly concentration-dependent; conditions with 5 mM metabolite supplementation comprise all cases with growth improvements, and 12 of the 15 cases with complete growth inhibition. Due to limitations in coverage of metabolites using mass spectrometry, we were unable to measure the consumption of urea, trimethylamine, methanol, ammonia, and acetate. However, PCC 6803 consumed 15 of the remaining 27 metabolites, with mixotrophy here defined as a statistically significant (p < 0.05, Welch’s t-test) reduction of at least 10% in signal intensity in either the 5 mM or 100 µM supplementation condition **(Fig. 2A).** Mixotrophy and final yield appear to be uncoupled, as exemplified in ten cases where PCC 6803 reached a lower yield (OD_750_ after ten days of growth) even when the metabolite was consumed, and three cases where PCC 6803 reached a higher yield when no metabolite was consumed. While these results demonstrate the breadth of mixotrophic capabilities of PCC 6803, they also illustrate that growth limitations cannot solely be attributed to carbon or energy limitations and hint that exogenous nutrients may interfere with other physiological processes.

Of the seven metabolites that improved growth, an increase in cell density above 10% after ten days of growth was observed with acetate, glucose, proline, and taurine. This reflects previous reports that PCC 6803 uses glucose and acetate for mixotrophic growth, increasing its yield[11]. ^14^C isotope experiments have demonstrated proline incorporation into PCC 6803 biomass in small quantities, however, whether it may be used in large quantities as a source of carbon or nitrogen is unknown[38]. Proline has also been recognized as an antioxidant whose intracellular concentration increases in cyanobacteria under high light conditions, and has been shown to have positive effects against a large number of environmental stressors in higher plants, although its mechanisms are not entirely understood[39–41]. Similarly, taurine has antioxidant properties against reactive oxygen species (ROS) in higher plants[42] which are also produced during cyanobacterial photosynthesis[43]. Additionally, taurine represents a potential source of sulphur that can be used to fuel the production of glutathione, a vital metabolite for the mitigation of oxidative stress in PCC 6803[44]. Proline was partially consumed when supplied at 100 μM concentrations while taurine was not, and neither proline nor taurine were consumed when supplemented at 5 mM **(Fig. 2A)**. These results suggest that the growth benefits of these metabolites may stem from their roles as antioxidants, not supplemental sources of carbon or nitrogen.

Supplementation of citrate, fructose, glutamine, glutarate, glutamate, succinate, malate, methionine, propionate, and threonine at 5mM all resulted in no growth, while lysine, propionate and threonine prevented growth even at 100 µM. The metabolites citrate, glutamine, glutamate, malate, and succinate are all involved in the tricarboxylic acid (TCA) cycle or are close derivatives of TCA intermediates **(Fig. 2B)**. Similar to many other cyanobacteria, PCC 6803 utilizes an incomplete TCA cycle characterized by the lack of 2-oxoglutarate dehydrogenase[45]. Metabolic flux analysis of PCC 6803 indicates a bifurcated topology within the TCA cycle enzymes that does not necessitate cyclic flow, and predicts that TCA activity must be coupled to growth[46]. As such, we hypothesize that supplementation of TCA cycle-related metabolites may impair TCA cycle function and have detrimental growth consequences.

### Condition-specific metabolite exudation

We performed untargeted metabolomics analysis on the cell-free culture supernatants to determine how the exudation of PCC 6803 changes in response to metabolite supplementation. Overall, we detected 925 ions, of which 62 could be annotated based on exact mass and were produced in at least one culture after ten days of growth. We then normalized each metabolite’s intensity across all conditions, allowing us to compare changes in the relative concentrations of each metabolite **(Supplemental Dataset 1).**

We identified which metabolites are produced most frequently by summing the normalized intensities of each metabolite across all conditions and ranking each metabolite according to its summed normalized intensity. Of the ten metabolites with the highest rank, five have the capacity to mitigate oxidative stress via reactive oxygen species (ROS) scavenging **(Fig. 3),** including dihydroneopterin, glutathione, formylglutathione, lactoylglutathione, and β-cyanoalanine[47–49]. Additionally, hydroxy-formykynurenine, the most frequently produced metabolite (produced in 44 of the 50 cultures with a visible OD_750_), is one of several compounds that cyanobacteria and other photoautotrophic organisms produce upon oxidative damage to photosystem II (PSII)[50,51]. These results suggest that many of our cultures may have experienced high light stress, and that the formation of, and metabolic response to, ROS was an important factor for the growth of PCC 6803 under our cultivation conditions.

**Figure 3:**
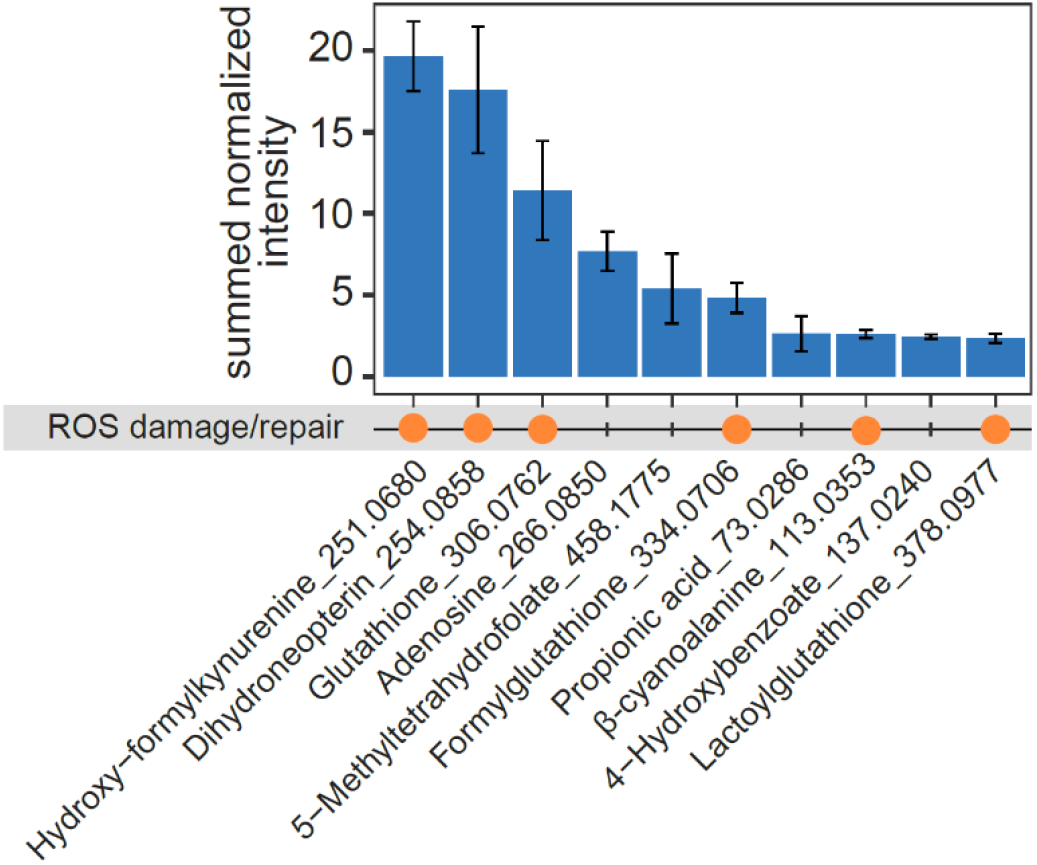
Most frequently produced metabolites across all conditions. Ten metabolites with the highest summed normalized intensity. Orange dots signify that these metabolites are involved in reactive oxygen species (ROS) damage/repair. For each annotated metabolite, the mass to charge ratio (m/z) is noted. All experiments are performed in triplicate, and error bars represent standard deviation around the mean.

We next examined changes in exuded metabolites, collectively referred to as the exometabolome, specific to each supplemented metabolite. To this end, we performed principal component analysis (PCA) on the exometabolome profiles of PCC 6803 in each growth condition **(Fig. 4A, Fig. S1).** The PCA revealed a clear separation of glucose and valine (supplemented at 5mM) from the other culture conditions in opposite directions along PC2. Glucose and valine exerted positive and negative influences on the growth of PCC 6803, respectively **(Fig. 4B)**, suggesting that differences in the exometabolome profiles are due to distinct underlying physiological responses. To investigate these responses, we identified which metabolites contribute to the separation observed in the PCA. Of metabolites that have the most positive or negative loading on PC1, most failed to show a clear separation between the valine and glucose conditions **(Fig. S2, S3)**. However, glucose and valine exometabolome profiles were clearly distinguished along PC2; therefore, we identified five metabolites with the most positive or negative loading on this component **(Fig. 4CD).** Gluconic acid, 5-methyltetrahydrofolate, formylglutathione, 2-phosphoglycolate, and itaconate, which have the most negative loading, were produced at high relative abundances under 5mM glucose supplementation and at higher concentrations than any other condition **(Fig. S4)**. The five metabolites exhibiting the most positive loading on the second principal component are produced under 5mM valine supplementation and no other condition: (iso)propylmalate, ketovaline, hydroxyglutarate, (Iso)leucine, and glutarate **(Fig. 4D)**.

**Figure 4:**
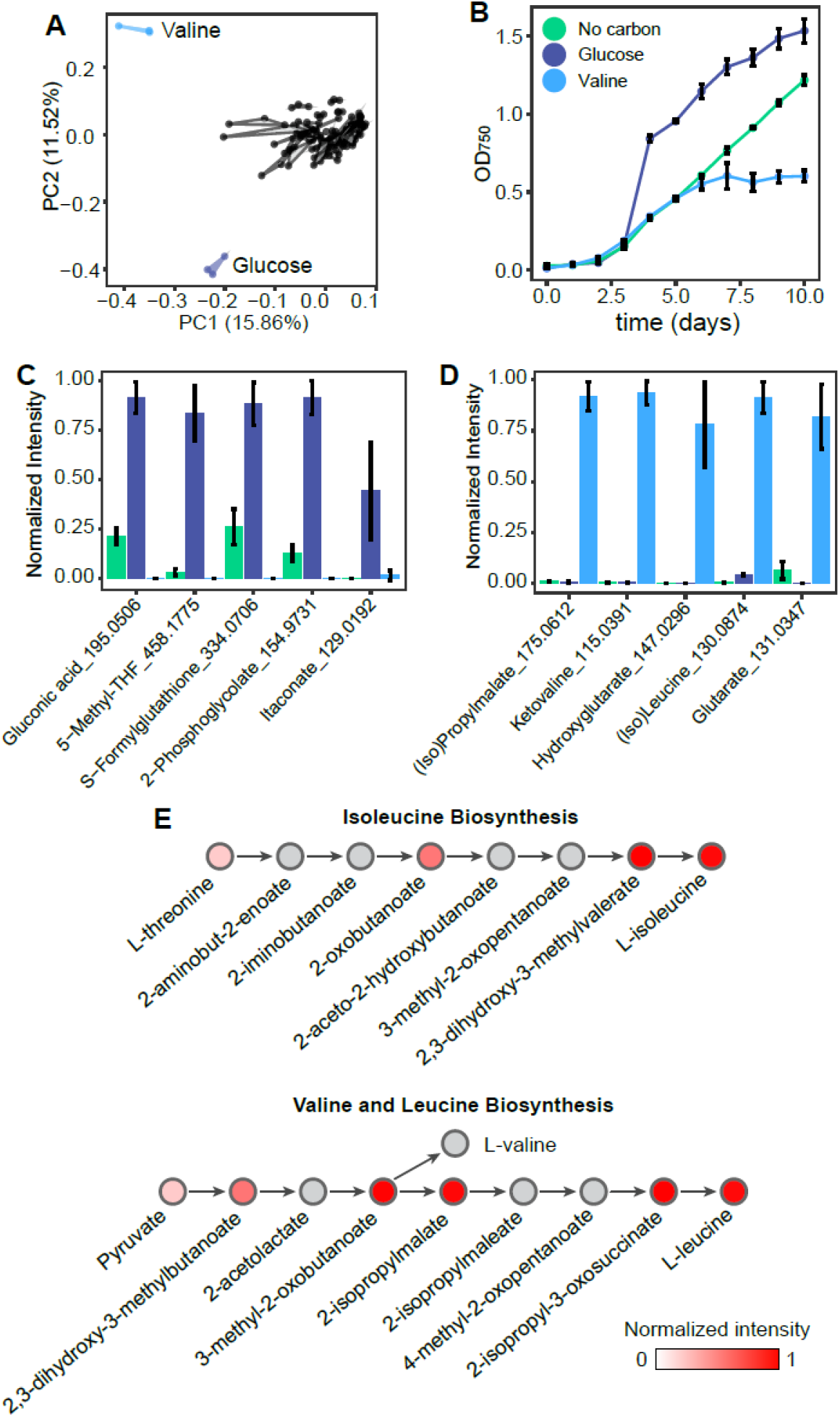
Condition-specific metabolite exudation. A) Principal component analysis of all exometabolite profiles in all growth conditions. B) Growth curve of PCC 6803 with or without 5 mM supplementation of glucose or valine. Metabolites with the C) most negative or D) most positive loading on the second principal component (PC2), and their normalized intensities for PCC 6803 growth with 5mM glucose (dark blue), 5mM valine (light blue) or no supplementation (green). E) Isoleucine, valine, and leucine biosynthesis pathways. The colour of the circle represents normalized intensities of metabolites produced with 5mM valine supplementation. Grey circles were not measured in our metabolomics dataset. Experiments were performed in triplicate and error bars represent standard deviation of the mean. Abbreviation: 5-methyltetrahydrofolate; 6-Methyl-THF

The exometabolome profile observed under valine supplementation suggests perturbations to amino acid metabolism, as all five metabolites that contribute to the separation of valine in the PCA are involved in either amino acid biosynthesis or degradation pathways. (Iso)propylmalate, ketovaline, and (iso)leucine participate in branched-chain amino acid biosynthesis, while hydroxyglutarate and glutarate are degradation products of lysine and/or glutamate. In addition, valine supplementation, similar to other tested amino acids, inhibits the growth of PCC 6803 **(Fig. 2A, 4B)**. Amino acid toxicity has been observed in PCC 6803 and other cyanobacteria, as well as chemoautotrophic and methylotrophic bacteria, and has been attributed to a stoichiometric imbalance in amino acid metabolism[9,52–54]. Another potential mechanism of growth inhibition may be due to altered intracellular carbon to nitrogen (C:N) ratios, which must be tightly regulated to maintain homeostasis[55].

To further explore the effects of valine supplementation on PCC 6803, we performed a metabolic pathway ranking to identify which pathways are most represented by the exuded metabolites (see methods). Branched chain amino acid biosynthesis is the highest ranked **(Table S1)**, as we measured eleven metabolites exuded after valine supplementation that are involved in this metabolic pathway **(Fig. 4E)**. These findings provide evidence for the hypothesis that the reduced growth conferred by high valine concentrations, and the resulting exudation of these metabolites, may be an adaptation to maintain optimal intracellular concentrations of amino acids or to maintain an optimal C:N ratio.

## Discussion

Despite the critical role of cyanobacteria as primary producers across different ecosystems, we lack a comprehensive analysis gauging both the growth and exudation response to a broad range of metabolites. Using a high-throughput cultivation platform, we were able assess the responses of *Synechocystis sp.* PCC 6803 to a large panel of supplemented metabolites, uncover how these metabolites change growth and exudation behaviour, and identify previously unknown metabolites potentially used for mixotrophic growth.

We screened through 32 metabolites at two concentrations and found that at least 15 metabolites were consumed mixotrophically. This includes six previously unreported metabolites: sucrose, malate, citrate, gluconate, methionine, valine. Moreover, our measurements of metabolites that are drawn down in the growth medium are mostly consistent with previous reports of ^14^C metabolite uptake. Two inconsistencies are alanine and lysine, which were not consumed in our experiments, but have previously been shown to be incorporated into PCC 6803 biomass[38,56]. Here, we find that lysine causes complete growth inhibition, even at 100 μM concentrations. Our results suggest that although PCC 6803 may encode active transporters for a given metabolite, in this case lysine, this metabolite may otherwise be inhibitory and is not used mixotrophically.

Supplementation most often inhibited, rather than enhanced, cyanobacterial growth. In particular, supplementation with TCA cycle intermediates or closely related compounds provided a high frequency of partial or complete growth inhibition at 5 mM concentrations. Disruptions to the bifurcated TCA cycle of PCC 6803 could stem from several sources. In PCC 6803, phosphoenolpyruvate from autotrophic carbon fixation enters the TCA cycle via conversion to oxaloacetate. The malate dehydrogenase enzyme in PCC 6803 has been shown to directionally favor the conversion of oxaloacetate to malate, rather than the oxidative direction of malate to oxaloacetate, at ∼361:1 [57]. Threonine, lysine and methionine are all synthesized from oxaloacetate. These compounds may represent a downstream sink for oxaloacetate, and supplementation may induce regulatory changes or product inhibition that necessitates the redirection of carbon flux to succinate, which would eventually accumulate or need to be assimilated via alternative pathways[58]. Similarly, glutamine is normally produced from 2-oxoglutarate and may represent a sink for 2-oxoglutarate. The presence of exogenously added glutamine would also likely disrupt this sink, resulting in the stagnation of flow of this branch of the TCA cycle. These metabolites may also directly interfere with the activity of TCA cycle enzymes. For example, citrate and succinate have been shown to cause significant reductions in fumarase activity[59], while citrate and malate reduce the activity of phosphoenolpyruvate carboxylase[60]. Meanwhile, high concentrations of propionate are toxic to PCC 6803 and other cyanobacteria[61], possibly via shrinking of the intracellular acetyl-CoA pool[62]. Collectively, the supplementation of central metabolites may cause disruptions to the TCA cycle and thus inhibit the growth of PCC 6803.

In most of our growth conditions, even without metabolite supplementation, PCC 6803 produced hydroxy-formykynurenine, a metabolite formed as a consequence to oxidative damage in PSII[50,51]. Complementary to this observation, five out of the ten most frequently produced metabolites are engaged in ROS scavenging, indicating an adaptive response of PCC 6803 to counteract oxidative stress. Together, these findings suggest that oxidative stress was a likely factor limiting growth. Though our cultures experienced a 12:12 light:dark cycle, all samples were collected in the light part of the cycle (approximately 4-6 hours after ‘sunrise’). Sampling throughout the light:dark cycle could clarify whether light stress contributed to the production of ROS-related metabolites. Oxidative stress occurs as a normal part of photosynthesis, particularly if light-driven electron transport proceeds faster than CO_2_ fixation, which consumes these electrons[43]. Concurrently, we found that supplementation at 5 mM of two metabolites with known antioxidant properties, proline and taurine [39,40,42,63], significantly improved growth. This suggests that ROS damage and repair are growth-limiting factors for PCC 6803, which may be alleviated by exogenous compounds with antioxidant properties.

We explored how metabolite supplementation changes the exudation behaviour of PCC 6803, and identified two conditions, glucose and valine supplied at 5 mM, where the exometabolome diverged compared to the other conditions. One metabolite exuded during glucose supplementation is gluconic acid, which can be formed from the direct oxidation of glucose or via the spontaneous hydrolysis of the oxidative pentose phosphate pathway (oPPP) intermediate, glucono-1,5-lactone. PCC 6803 relies on the oPPP as a high-flux route for sugar catabolism[64], meaning that gluconic acid may be produced as an overflow product or unwanted side product as a consequence [64,65]. A similar observation has been made in the Alphaproteobacterium *Gluconobacter oxydans*, wherein high glucose concentrations not only induce gluconic acid production, but also prevent its uptake[66]. Additionally, 2-phosphoglycolate is produced by ribulose 1,5-bisphosphate carboxylase (RuBisCo) as an unwanted by-product of photorespiration, where oxygen competes with carbon dioxide for the desired carboxylation reaction in photosynthesis[67]. Because glucose increases the yield of PCC 6803, which will inherently have a higher demand for CO_2_, increased photorespiration and 2-phosphoglycolate concentrations under glucose supplementation may be a consequence of a depleted CO_2_ supply. However, it is unclear why glucose supplementation resulted in a more divergent exometabolome profile than other metabolites that also significantly improved growth, such as acetate, asparagine, and gluconate.

In contrast to glucose, valine significantly impaired the growth of PCC 6803 when supplied at 5 mM. We identified that the exuded metabolites overwhelmingly represented pathways including branched chain amino acids and their precursors, as well as several other amino acid degradation products. The exudation of these metabolites might indicate an attempt by PCC 6803 to restore its intracellular amino acid balance or C:N ratio. However, supplementation of other branched chain amino acids did not have the same effect – in fact, (iso)leucine supplementation at 5 mM significantly improved the growth of PCC 6803, although it was not consumed. Other amino acids also did not affect the exometabolome in this way, although many were consumed to a similar degree and also negatively affected growth. One explanation for this observation could be that valine alters the regulation of branched chain amino acid biosynthesis, as it has been shown in other organisms that valine specifically causes a feedback inhibition on acetohydroxyacid synthase[68]. However, further investigation of the regulatory influence of valine in PCC 6803 must be performed to better understand this observation.

Using high-throughput metabolomics, we were able to measure the consumption and exudation of low molecular weight, polar metabolites including amino acids, sugars, and organic acids; however, we did not measure extracellular polymeric substances (EPS), which are an important high molecular weight component of cyanobacterial exudates[20]. EPS can also provide carbon to heterotrophic bacterial populations[13,20,69] and are important for the colony-forming lifestyles of certain cyanobacterial strains[70]. Recent work has shown that the composition of PCC 6803 EPS changes with environmental nutrient conditions[71,72], but to our knowledge no screen like that presented here has been conducted with EPS quantity or composition as a readout. Such work will be important for completing our understanding of ecologically relevant cyanobacterial responses to diverse nutrient conditions.

With our high-throughput culturing platform, we were able to provide insight into how cyanobacterial physiology and metabolism responds to a large range of metabolites. This platform could be leveraged for other applications involving the cultivation of photoautotrophs under a large set of conditions, expanding the possibilities of research questions and feasible experimental setups. This could involve fields such as toxicology, where it is of interest to understand how cyanobacteria and algae respond to an ever-growing range of xenobiotics and other environmental pollutants, or microbial ecology, where the interactions among photoautotrophs and other microbes could be studied in a high-throughput manner.

## Materials and Methods

### Chemicals and culturing conditions

Axenic PCC 6803 was obtained from the Pasteur Culture Collection (Paris, France) and precultures were maintained in 100 mL flasks by transferring to fresh media weekly. The medium for precultures was BG-11 medium (Sigma) supplemented with 25 mM HEPES sodium salt, adjusted to pH 7.5. For the screen, we used BG-11 composed of three stock solutions, supplemented with trace metals and sodium nitrate, with the addition of 25 mM HEPES sodium salt. The three stock solutions were prepared as follows: 0.1 g/L magnesium disodium EDTA, 0.6 g/L ferric ammonium citrate, 0.6 g/L citric acid monohydrate, 3.6 g/L calcium chloride dihydrate (stock solution 1); 7.5 g/L magnesium sulphate heptahydrate (stock solution 2); 3.05 g/L potassium phosphate anhydrous (stock solution 3). The trace metal solution was made as follows: 2.86 g/L boric acid, 1.81 g/L manganese chloride tetrahydrate, 0.22 g/L zinc sulphate heptahydrate, 0.05 g/L copper sulphate anhydrous, 0.05 g/L cobalt(II) chloride hexahydrate, 0.39 g/L sodium molybdate dihydrate. The stocks were combined as follows: for 1 liter of medium, 10 mL each stocks 1-3, 1 mL trace metals stock, 18 mL 1M sodium nitrate stock, 100 mL 250 mM HEPES, and 851 mL sterile ddH_2_O. The stock, trace metals, HEPES and sodium nitrate solutions were all filter sterilized (0.22 µM) and stored at 4°C prior to use. BG-11 media was made fresh for each preculture transfer or experiment. 0.5 M stock solutions of all supplemented metabolites were prepared in ddH_2_O, and filter sterilized (0.22 µM). The stocks were stored at 4°C and diluted accordingly into fresh BG-11 media for each experiment. Unless otherwise noted, all chemicals were obtained from Sigma Aldrich. All growth experiments were conducted at 30 °C, with a light intensity of 70 µmol m^-2^ s^-1^ supplied by 4 neutral white LED modules (Waltron) and a 12:12 light:dark cycle. Cultures were continuously shaken at 220 rpm (100 mL flasks) or 400 rpm (24-well plates) with an orbital shaker (VWR) placed inside the incubator. Cyanobacterial growth was quantified by measuring optical density at 750 nm (OD_750_) using a Tecan Infinite Nano microplate reader (Tecan). Cultures in 24-well plates were inoculated from precultures to an OD_750_ of 0.01, and with a starting volume of 1.6 mL. To account for evaporation (approximately 60 μL per day per well, data not shown) and loss of volume due to metabolomics sampling, we replenished all wells with 240 μL water on day 4, and 180 μL on day 7. We also replenished the 100 μL that was sampled for metabolomics on day 5.

### Vented lids

To enable robust photoautotrophic growth in 24-well plates, custom-made vented lids were used to increase aeration in the wells. To make the lids, holes were made in the space between the condensation rings of a polystyrene 24-well plate lid (Corning) with an 18G hypodermic needle heated using a Bunsen burner. The holes were covered by attaching a square of PVDF filtration membrane (0.45 µm pore size, Millipore) on the inner face of the lid using clear PVA glue (UHU) to prevent airborne contamination during cultivation. The holes were positioned between, rather than on top of, the wells to avoid shading the cultures from the overhead lights in the incubator. Before use, the lids were wrapped in a single layer of cling film and sterilized by exposure to 254 nm UV radiation for 15 minutes inside a Spectrolinker XL-1500 UV-crosslinker (Spectro-UV). The lids were then placed onto pre-sterile 24-well plates in a sterile hood after plates had been inoculated with cultures. The edges of the plates were sealed with Parafilm to prevent excess evaporation.

### Exometabolomics

For sample collection, 100 µL of culture was added to a 96-well v-bottom plate and centrifuged at 2500rcf for 20 minutes. The supernatant was collected and stored at –20 C° until measurement. The sample was diluted 10-fold and measured using LC-QTOF-MS. Measurements were performed using an Agilent 6520 Time of Flight Quadrupole Time of Flight Mass Spectrometer in negative mode, high resolution mode with a 0.9 Hz scan rate and an acquisition mass range of 50-1700m/z. The drying gas was set to 10 L/min, nebulizer to 30 psig, and gas temperature to 325 C°. Using an Agilent 1100 series liquid chromatography stack, 3 µL of sample was injected into an Agilent EC-CN Poroshell column (2.7 µm, 50 x 2.1mm) using an adapted salt tolerant method to mitigate ion suppression caused by salts in the medium[75]. The column was kept at 20°C and the flow rate was 350 µL/min. The buffer used was 10% Acetonitrile (CHROMASOLV), 90% mass spectrometry grade water, and 0.01% Formic acid. The method was operated isocratically, and every 40 samples the column was washed by flushing for five minutes with 90% Acetonitrile (CHROMASOLV), 10% mass spectrometry grade water, and 0.01% Formic acid, followed by equilibration for ten minutes with the isocratic buffer. Raw data for all measurements was subjected to a spectral processing and alignment pipeline using Matlab (The Mathworks, Natick) as described previously[76]. Peaks were annotated based on exact mass by matching them to metabolites predicted to partake in the PCC 6803 metabolic network according to the Biocyc Database[77]. Raw spectral files have been deposited to the MassIVE database under accession MSV000093130 with password reviewer123, and will be made publicly available upon acceptance of this manuscript.

To compare metabolite production between culture conditions, all produced metabolites were normalized between 0 and 1 (supplemental dataset 1). First, we calculated a limit of detection for each metabolite. This is the average metabolite intensity for BG-11 medium background sample, plus three times standard deviation of these background intensities. Any metabolite intensity that was below the limit of detection was reassigned to have the same intensity as the background medium. To remove the influence of supplemented metabolites on the normalization, for samples that have a supplemented metabolite, the intensity of this metabolite was reassigned to have the same intensity as the background medium. Next, all metabolite intensities were normalized between 0 and 1, where 0 is the intensity of the background medium and 1 is the intensity of the sample that has the highest concentration of the metabolite.

### Metabolic pathway ranking

Using all annotated metabolites that were detected in any culture condition, we identified which metabolic pathways they participate in based on the Biocyc pathway database of PCC 6803. This yielded the total number of detected metabolites for each metabolic pathway. We then calculated the number of metabolites found in each metabolic pathway separately for each of our 65 culture conditions. Then, we scored each metabolic pathway’s importance in each culture condition by dividing the number of produced metabolites in the pathway by the total of detected metabolites across all pathways in that condition. We then ranked the pathways by this score to identify which pathways are most represented by the metabolites exuded in each culture condition.

## Acknowledgements

We gratefully acknowledge financial support from the Simons Foundation through the Principles of Microbial Ecosystems (PriME) collaboration (grant 542395) and the ETH Career Seed Award. ZL was supported by HFSP (LT000192/2018-L).

## Supplementary information

**Supplementary table 1:**
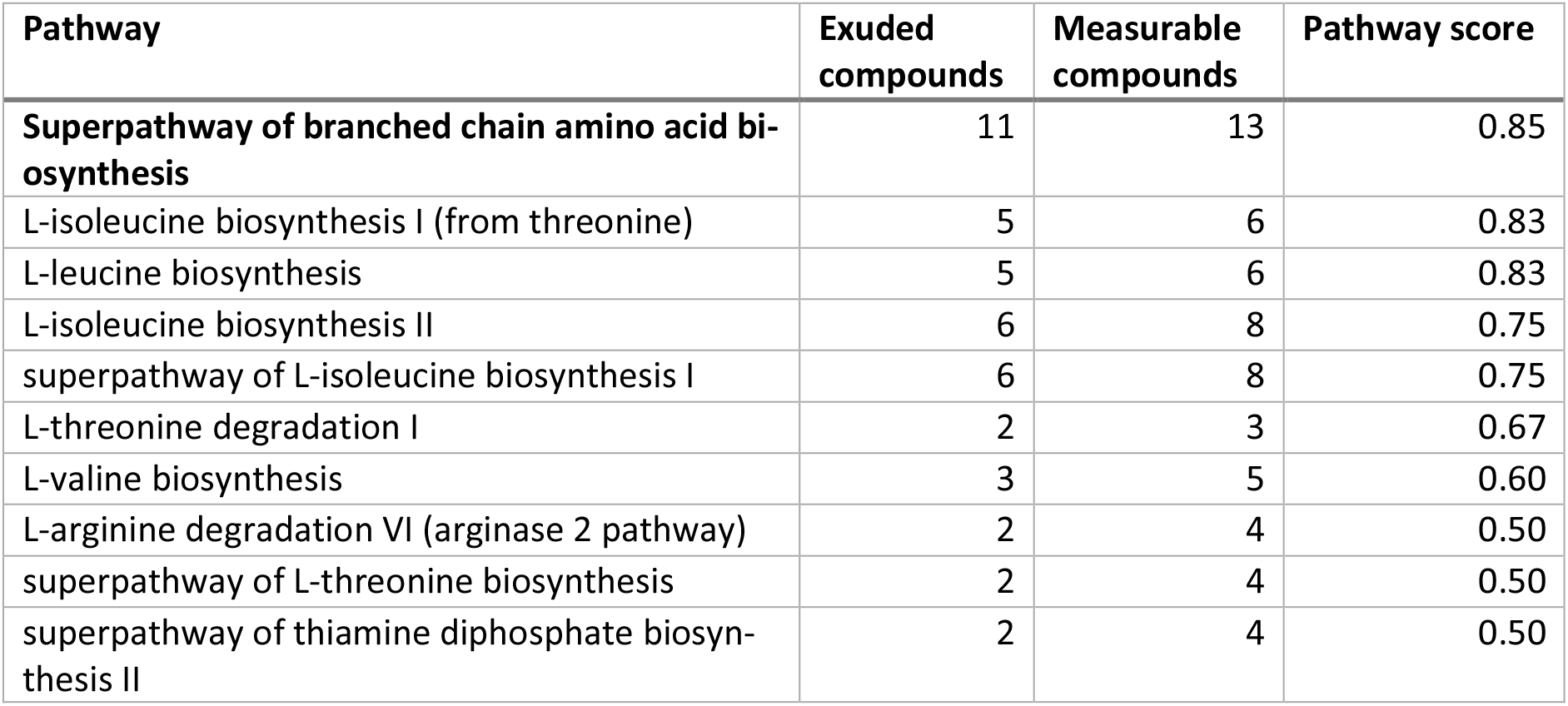
Metabolic pathways with the highest rank when PCC 6803 is grown with 5 mM valine, based on the exometabolome profile. Measurable compounds are the total number of compounds in the pathway that were measurable in any growth condition. Exuded compounds are the number of compounds in the pathway that were exuded in the 5 mM valine condition. The pathway score is the percentage of measurable compounds that were exuded, and all pathways were ranked based on this score.

**Supplementary figure 1:**
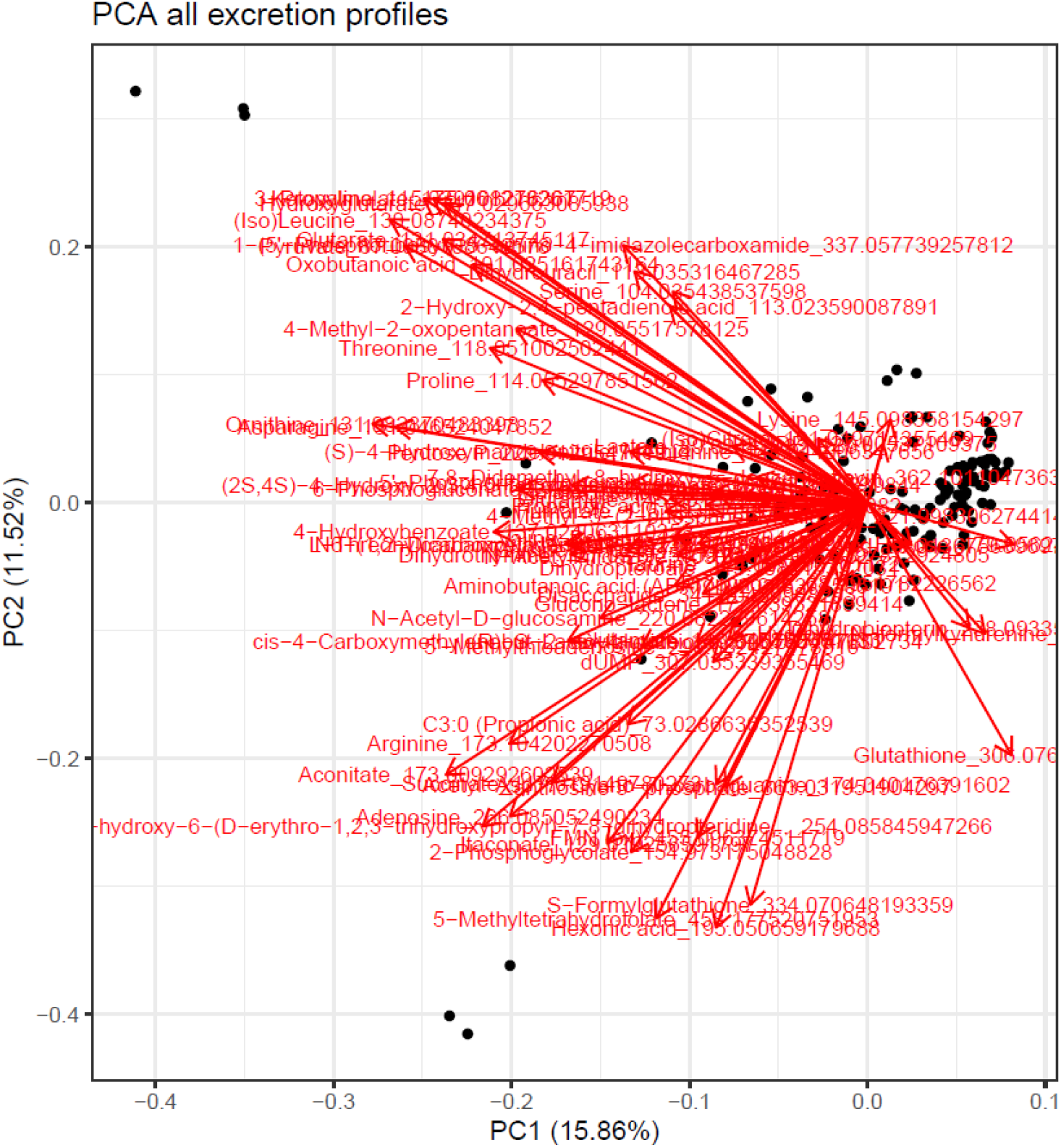
PCA loading plot of all secreted metabolite profiles.

**Supplementary figure 2:**
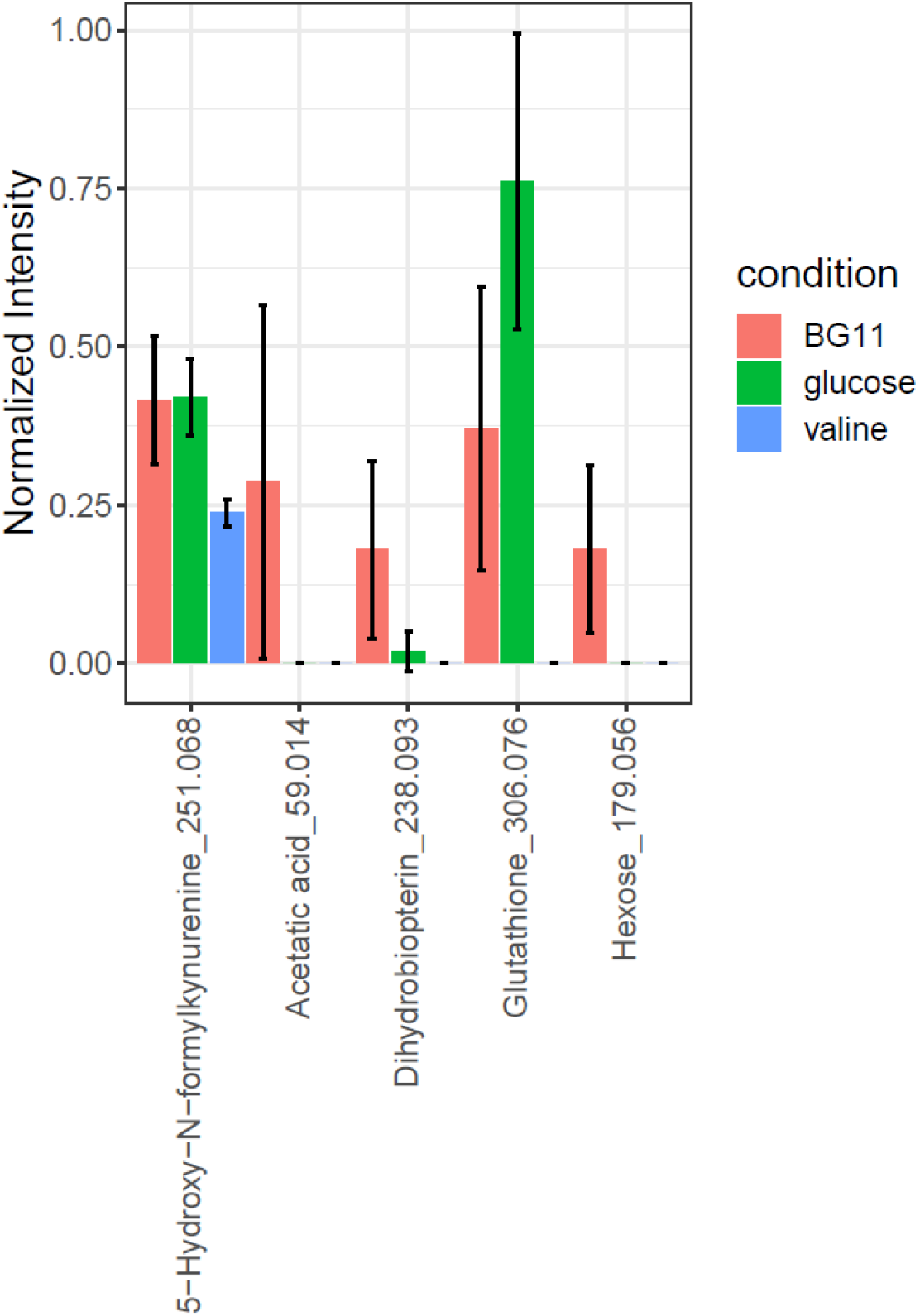
Normalized intensity of metabolites that have the most positive loading on the first principal component. Error bars represent standard deviation of the mean.

**Supplementary figure 3:**
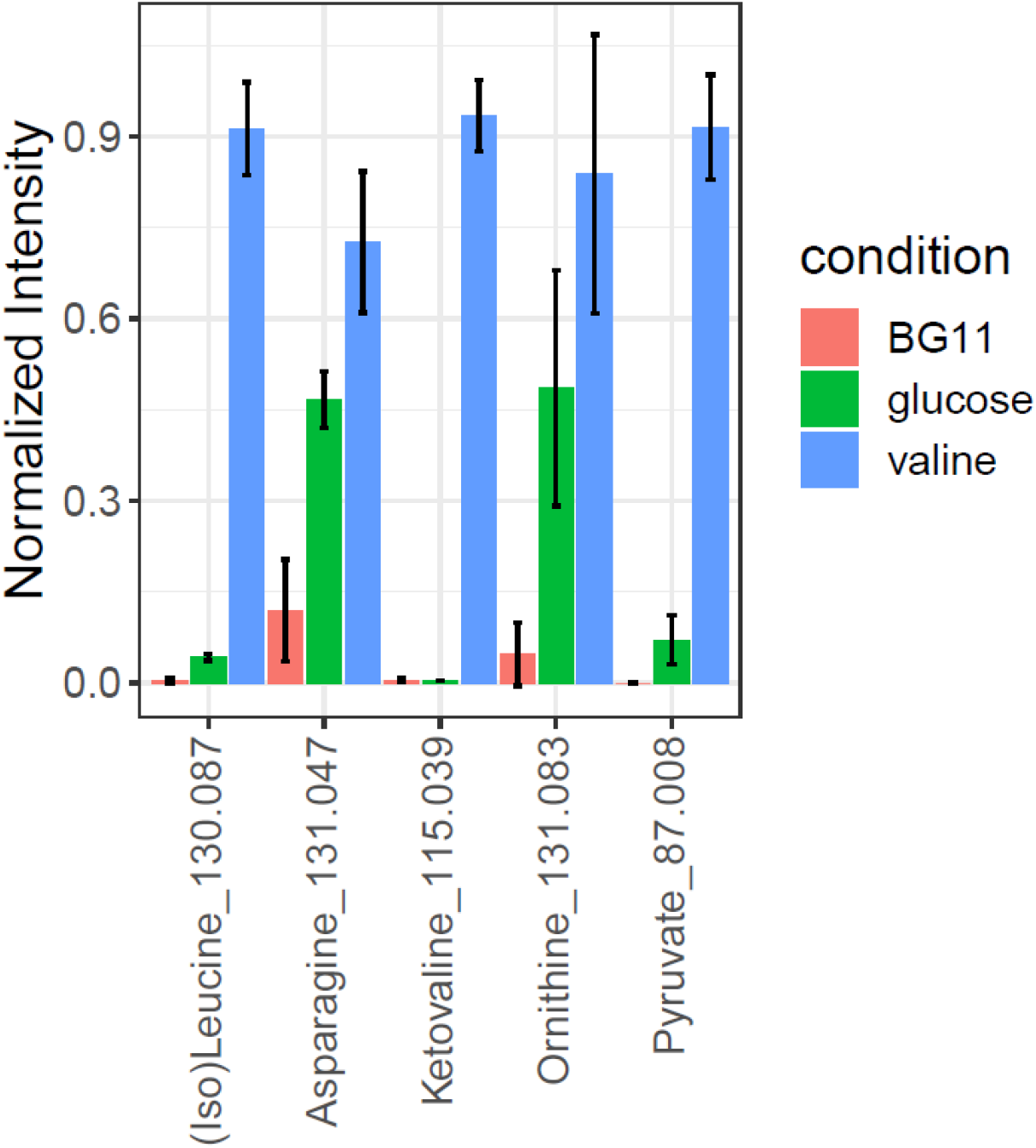
Normalized intensity of metabolites that have the most negative loading on the first principal component. Error bars represent standard deviation of the mean.

**Supplementary figure 4:**
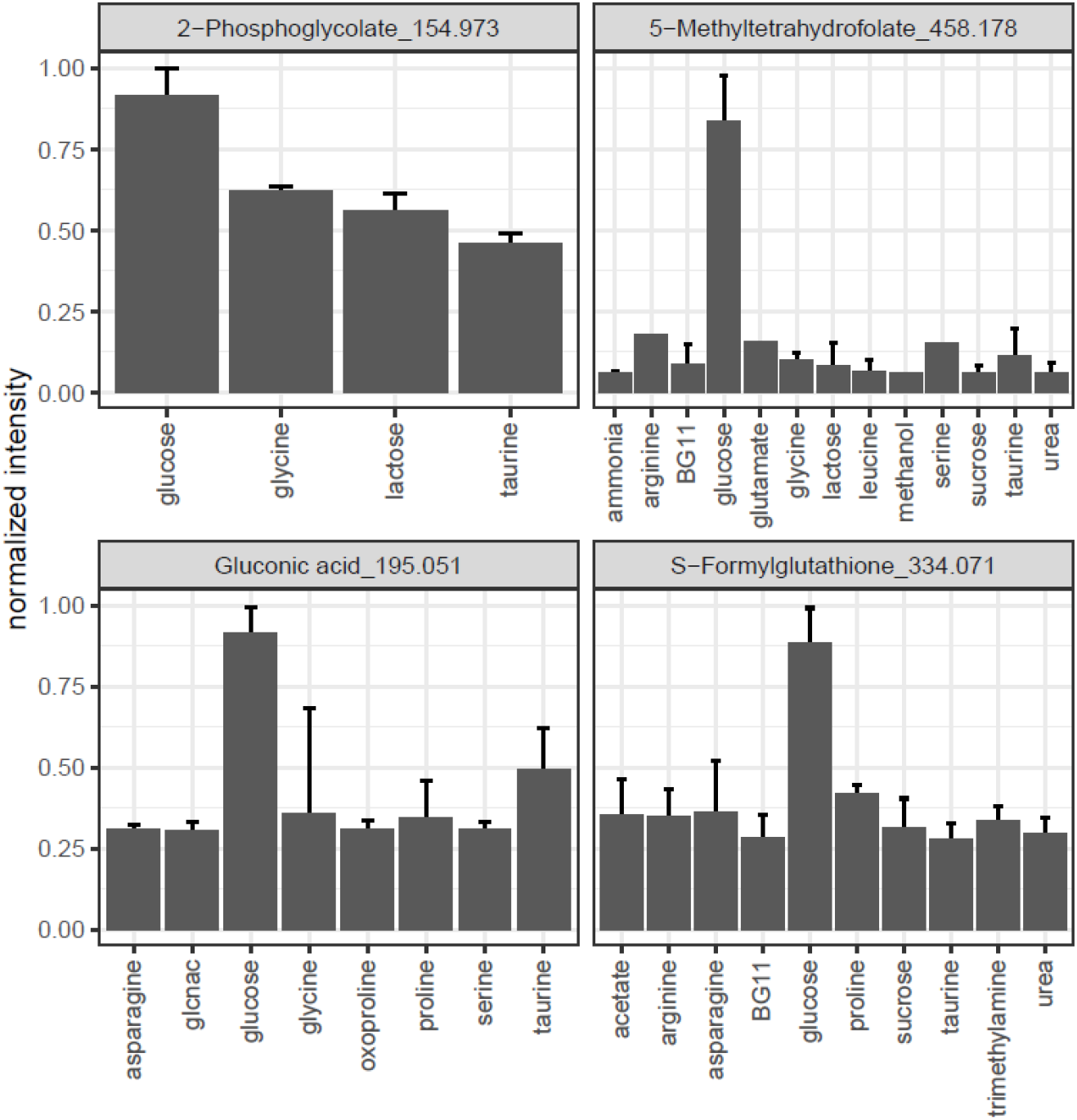
Metabolite concentrations of gluconic acid, S-Formylglutathione, 2-Phosphoglycolate and 5-methyltetrahydrofolate, which contribute most to the negative loading on the second component of PCA. All conditions in which the metabolite is detected above the limit of detection are listed. Not shown is itaconic acid, which is only detected with 5mM glucose supplementation. Error bars represent standard deviation of the mean of three replicate cultures.

